# Metagenomic analysis and keeping quality determination of stored pepper pastes

**DOI:** 10.1101/2025.03.14.643245

**Authors:** A. O. Fawole, K. Ogungbemi, G. O. Onipede, A. L. Kolapo, E. O. Okedara

## Abstract

Pepper paste is a popular condiment used in many cuisines around the world. However, stored pepper paste can deteriorate due to microbial growth and other factors. This study used metagenomic analysis to study the microbial communities in stored pepper pastes produced in a cottage setting, with or without onions labelled ‘A’ and ‘B’, respectively. Microbial count and chemical analysis were also used to determine their keeping quality. The bacterial count in the pepper pastes increased over time. While there was no yeast growth in sample ‘A’, the yeast number in sample ‘B’ increased from 3.0 to 7.0 Log CFU as the day of storage increased. In the selective culturing, there was no *Staphylococcus aureus* in both samples, growth of enteric bacteria was found in both products towards the end of the storage period, and *Salmonella species* was only found at some point in ‘B’ (3.0 Log CFU). *Proteus* (51%) and *Pediococcus* (48%) were the highest competing genera in ‘A’, while *Bacillus* (84%) dominated ‘B’ microbiota. The acidity of the samples increased while the nutrients became depleted as storage day increased. At the end of the study, none of the samples without onions remained physically stable. Regardless of any visible deterioration, this study recommends consuming pepper with onions within 60 days and pepper without onions within 30 days.

## Introduction

Culturing is a valuable method for identifying and quantifying microbes in food. It involves techniques like plate counting and selective culturing, which can estimate the number of viable microbes and encourage the growth of certain types of microbes while inhibiting others. However, culturing has limitations as it can only identify microbes that can be grown in the chosen medium, and some microbes may not grow at all. Despite these limitations, it remains an essential tool for assessing the safety of food (Chauhan & Jindal, 2020).

Methods for identifying microbiota in food without relying on culture-based techniques are gaining popularity due to their ability to provide a complete understanding of microbial communities. DNA-based methods such as polymerase chain reaction (PCR) and next-generation sequencing (NGS) are frequently used in non-culture-based analyses. These methods can potentially revolutionise our comprehension of food microbiota and their impact on safety and quality (Quigley et al., 2012). Metagenomic analysis is a potent tool for studying microbial communities in various environments, including food. It involves direct analysis of genetic material from environmental samples, bypassing the need for traditional culture-based methods. This offers a more comprehensive view of microbial diversity and dynamics in food and their potential effects on safety and quality. Recent studies have shown that metagenomic analysis can identify foodborne pathogens and spoilage-related microbes and monitor microbiota during food production and storage (Etebu et al., 2018).

The microbiota in pepper components and their products vary depending on intrinsic and extrinsic factors. Pepper paste is a popular ingredient in many countries. It could refer to a paste made using one type of pepper, like bell pepper or cayenne pepper, or it could be a blend of ingredients such as tomato, bell pepper, scotch bonnet chilli, cayenne pepper, and onions. The quantity of each component used in a blend varies depending on the chef’s preference, the company recipe, and the customers’ preference in a business setting. Additionally, different varieties of the components are mixed based on availability and preferences. Since the components are seasonal, processors have devised means of preservation, such as refrigeration, salting, drying, processing into a paste, using modern equipment and methods to process, as well as using different packaging methods (Chitravathi et al., 2015; Aşkin, 2018).

Cayenne pepper is a non-climacteric fruit. It is highly perishable and tends to deteriorate quickly during postharvest handling and storage, which results in significant losses. Postharvest problems include quality degradation, chilling injury, and shrivelling due to rapid weight loss. In ambient conditions, it changes colour and deteriorates within a few days after harvest (Chitravathi et al., 2015). Tomato, bell pepper, and chilli-scotch bonnet have similar postharvest issues as cayenne pepper, according to researchers (Falola et al., 2023). Apart from reduced quality and quantity, pepper components become unaffordable when out of season.

To address the aforementioned issues, some African processors have opted to create large batches of mixed pepper paste in a cottage setting while the components are fresh and in season. This paste is packaged in repurposed mayonnaise bottles and stored at room temperature. According to local processors, households consume the stored pepper paste over a period of 9 to 12 months, provided there are no signs of spoilage. However, the literature has limited information regarding the quality and stability of pepper paste produced in this manner. This investigation aims to assess the chemical and microbiological characteristics of homemade pepper pastes that are not manufactured following any established national standards or industrial facilities. Through these observations, the study proposes an appropriate timeline for the pepper paste’s shelf-life. The experimental design was based on input from local producers from a wide range of Yoruba and Ibo communities in Nigeria. Two sets were created: one with onions and one without.

## Materials and methods

### Samples preparation

Varieties of fresh ripe red pepper: bell pepper (*Capsicum annuum*); chilli-scotch bonnet (*Capsicum chinence*); and cayenne pepper (*Capsicum annuum var. acuminatum*) with tomatoes (*Lycopersicon esculentum*) and onions (*Allium cepa*) were procured from a local market in Ibadan, Nigeria. The pepper components were sorted to remove defective ones and washed with tap water. Onions were peeled, washed, and cut into medium sizes of 3 cm ± 2 to ease grinding. They were two categories: All pepper components with onions (Pepper paste/Sample A), and pepper components without onions (Pepper paste/Sample B).

### Pepper paste treatment, packaging and storage

The pepper components were mixed in a ratio of 8:4:4:2:3 for tomatoes, bell pepper, cayenne pepper, chilli-scotch bonnet and onions (where applicable), respectively. Each category was ground separately using a locally fabricated grinding machine (Engine capacity 6.5 hp). The pastes produced after grinding were boiled at 90 °C ± 2 (Extech-SD200, 3-Channel temperature datalogger) for an hour to kill microorganisms that may be present and thicken the samples. Pre-washed and steamed glass jars (Twenty-four jars for each category) were filled with hot pastes to a marked maximum level and loosely corked because of the next stage of steaming the content for 1 min in boiling water. Then, the bottles were tightly corked and stored on clean, dust-free shelves at room temperature until when physical signs of spoilage were noticed. The method of processing, packaging and storing the ‘pepper paste A’ was from personal communication with three domestic producers, while category B was prepared for a control experiment.

### Microbiological examination of pepper pastes

The study conducted bacterial and fungal counts using Plate Count Agar (PCA) and Potato Dextrose Agar (PDA) respectively. The presence of specific bacteria was also checked using selective media such as Mannitol Salt Agar (MSA), Bismuth Sulphite Agar (BSA), and MacConkey Agar (MA). The analyses were repeated every 30 days until samples deteriorated.

### DNA extraction and metagenomic analysis of pepper pastes

DNA extraction and metagenomics analysis were carried out as a modified procedure to the one described by Etebu *et al*. (2018). DNA was extracted and purified from pastes on the 60th day of production. The sample was lysed by bead beating and centrifuged, and nucleic acids were collected and transferred to a filter collection tube. The filtrate was mixed with DNA binding buffer, transferred to an IC column, and centrifuged twice. The DNA was washed twice with DNA pre-wash buffer and DNA wash buffer before being transferred to a clean micro- centrifuge tube and incubated with DNase/RNase-free water for 1 min. The DNA samples were thereafter sent to Inqaba Biotechnology Pretoria South Africa for metagenomic analysis of full length 16s gene amplicons. Samples were sequenced on the Sequel system by PacBio targeting the 16S rRNA gene variable region V3–V4 from the genomic DNA. Raw sub-reads were processed through the SMRTlink (v11.0) Circular Consensus Sequences (CCS) algorithm to produce highly accurate reads (>QV40). These highly accurate reads were then processed through vsearch (https://github.com/torognes/vsearch) and taxonomic information was determined based on QIMME2.

### Chemical analysis of pepper pastes

All parameters checked were done in triplicate.

### Determination of pH of pastes

The pH was measured on 5 g of each sample dissolved in 50 ml of distilled water. It was determined using a MW180 Max pH meter (pH/mV/EC/TDS/NaCl) equipped with an electrode (MA917B/1).

### Determination of ash content of pastes

The paste sample (3 g) was charred before being burned in a furnace (550 ºC for 8 h) to remove all organic material (Fawole, 2019). The remaining inorganic material produced white ash, which was weighed to determine the percentage ash content as stated in Eq. 2.1.

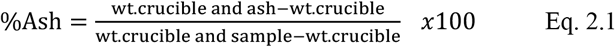

### Determination of titratable acidity (TA) of pastes

The samples (10 mg each) were mixed with distilled water (25 ml) and filtered before being titrated with 0.1 M NaOH and phenolphthalein (AOAC, 2000). The resulting citric acid percentage was used to express titratable acidity. Percentage TA was determined using Eq. 2.2 [V = volume of 0.1M NaOH, M = molarity of NaOH and F = factor of citric acid (0.007005)]:

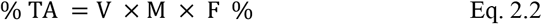

### Determination of dry matter of pepper pastes

The dry matter content of pepper pastes was determined by drying a sample in an oven and weighing it in 5-minute intervals until a constant weight was reached (AOAC, 2000). The percentage of dry matter was calculated using the weight of the crucible and sample before and after drying (Eq. 2.3).

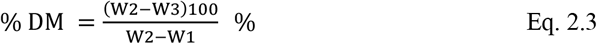

W1= crucible weight in g; W2= crucible-sample weight in g before oven drying; W3= crucible- sample weight in grams after oven drying

### Determination of total carotenoids of pastes

Total carotenoids (TC) in pepper paste were determined through a spectrophotometric method (AOAC 2000). A 12.5 ml of a solvent mixture (20 ml ethanol, 30 ml n-Hexane, and 2 ml 2% NaCl) was used to extract a 2.5 mg sample, which was then transferred to a separating funnel and allowed to stand for 10 min. The upper hexane layer was collected and its absorbance was measured at 436 nm using Spectrumlab 22PC10069. TC was calculated using the formula below:

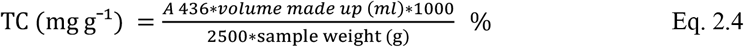

### Physical examination of pastes

The sample was inspected monthly for any signs of spoilage, including colour change, smell, and mould growth. Samples with no signs of spoilage were used for analysis while the ones with mould growth were discarded.

### Statistical Analysis

The data obtained for chemical analysis were analysed using the SPSS 22 software package; a One-way ANOVA was carried out to determine significant differences; and Tukey’s multiple range tests were used to separate means.

## Results and discussion

The quality, shelf-life and bacterial community of pepper pastes produced with and without onions were evaluated by microbiological, metagenomic and chemical analyses. The results showed the trends in quality depletion and the concern this could generate on safety.

### Microbiological examination of pastes

Bacteria and moulds were noticed on the 60^th^ day of storage (Table 1). Sample ‘A’ with onions had a 1.2 log increase in bacteria count, while sample ‘B’ without onions had a 3.0 log increase. Yeasts were not detected in ‘A’ but were present in ‘B’ from day zero, with an increasing log count as the storage days increased. Mould log counts increased in ‘A’ but decreased in ‘B’. By day 120, all remaining samples in ‘B’ had gone bad. The effect of adding onions to sample ‘A’ was evident in the observed trend of the microbial counts of the stored pepper pastes. Previous studies have extensively investigated the antimicrobial activity of crude onion, which could explain the low counts of bacteria and moulds found in ‘A’ (Santas et al., 2010; Kabrah et al., 2016). The absence of yeast in sample ‘A’ throughout the study further established onions’ absolute inhibitory activity against this group of microorganisms. Researchers have used different onion varieties, forms, and extraction techniques to control infectious yeasts and bacteria (Chakraborty et al., 2022; Oyawoye et al., 2022).

**Table 1.** Total viable count of Bacteria, Fungi, *Staphylococcus aureus, Salmonella species* and enteric bacteria within days of storage.

| Organism/Medium | Sample | Unit | Storage day(s) |  |  |  |  |
| --- | --- | --- | --- | --- | --- | --- | --- |
|  |  |  | 0 | 30 | 60 | 90 | 120 |
| Bacteria on PCA | A | CFU ml<sup>-1</sup> | ND | ND | 1.0 x 10<sup>3</sup> | 1.0 x 10<sup>4</sup> | 1.5 x 10<sup>4</sup> |
|  |  | Log CFU | - | - | 3.0 | 4.0 | 4.2 |
|  | B | CFU ml<sup>-1</sup> | ND | ND | 1.0 x 10<sup>4</sup> | 1.0 x 10<sup>7</sup> | NA |
|  |  | Log CFU | - | - | 4.0 | 7.0 | - |
| Yeast on PDA | A | CFU ml<sup>-1</sup> | ND | ND | ND | ND | ND |
|  |  | Log CFU | - | - | - | - | - |
|  | B | CFU ml<sup>-1</sup> | 1.0 x 10<sup>3</sup> | 1.4 x 10<sup>4</sup> | 1.0 x 10<sup>5</sup> | 1.0 x 10<sup>7</sup> | NA |
|  |  | Log CFU | 3.0 | 4.1 | 5.0 | 7.0 | - |
| Mould on PDA | A | CFU ml<sup>-1</sup> | ND | ND | 1.0 x 10<sup>2</sup> | 1.0 x 10<sup>3</sup> | 1.0 x 10<sup>3</sup> |
|  |  | Log CFU | - | - | 2.0 | 3.0 | 3.0 |
|  | B | CFU ml<sup>-1</sup> | ND | ND | 1.0 x 10<sup>5</sup> | 1.0 x 10<sup>3</sup> | NA |
|  |  | Log CFU | - | - | 5.0 | 3.0 | - |
| *Staphylococcus aureus* on MSA | A | CFU ml<sup>-1</sup> | ND | ND | ND | ND | ND |
|  |  | Log CFU | - | - | - | - | - |
|  | B | CFU ml<sup>-1</sup> | ND | ND | ND | ND | NA |
|  |  | Log CFU | - | - | - | - | - |
| *Salmonella species* on BSA | A | CFU ml<sup>-1</sup> | ND | ND | ND | ND | ND |
|  |  | Log CFU | - | - | - | - | - |
|  | B | CFU ml<sup>-1</sup> | ND | ND | 1.0 x 10<sup>3</sup> | 1.0 x 10<sup>3</sup> | NA |
|  |  | Log CFU | - | - | 3.0 | 3.0 | - |
| Enteric bacteria on MCA | A | CFU ml<sup>-1</sup> | ND | ND | ND | 2.4 x 10<sup>4</sup> | ND |
|  |  | Log CFU | - | - | - | 4.4 | - |
|  | B | CFU ml<sup>-1</sup> | ND | ND | 1.9 x 10<sup>7</sup> | 2.7 x 10<sup>3</sup> | NA |
|  |  | Log CFU | - | - | 7.3 | 3.4 | - |

Sample A- Pepper paste with onions
Sample B- Pepper paste without onions
ND- Not detected, NA- No analysis, as the copies of the sample were overtly spoilt

Sources of bacteria and fungi in the final products could be from raw materials, grinder, and glass bottles for packaging and handling of processors (Banwart, 1989). The contamination is possible since the study was carried out according to domestic processors who do not follow any established cottage food production guides but their cleaning instincts. The boiling temperature of pepper paste, steaming, and hot filling of the glass bottles are stages in the production that could have killed some microbes and suppressed the others that were made non-viable until after 30 days. Russell (2003) posited that many non-sporulating bacteria are quickly rendered inactive at about 50 °C and beyond, with the inactivation rate increasing as the temperature rises. On the contrary, bacterial and fungal spores are far more resilient, and significant reductions in viability typically require minimum moist heat temperatures of 100 °C and frequently significantly higher.

*Staphylococcus aureus* was not found in the products (Table 1). However, *Salmonella species* were detected in the sample without onions by day 60 (3.0 Log CFU). In sample A (with onions), the enteric count was 4.4 Log CFU on the 90^th^ day, while sample B (without onions) showed growth on the 60^th^ and 90^th^ days with 7.3 and 3.4 Log CFU, respectively. The enteric bacterium was not detected in ‘A’ on day 120 of the study, and a decrease in count from day 60 to day 90 in ‘B’ was recorded. The discrepancy in the record of enteric bacteria growth over the course of the study suggests contamination from the bottles of the copies of the samples that were analysed at the periods when growth was observed. As much as packaging is indispensable in food manufacturing, reusing or handling it inappropriately can lead to contamination (Lau and Wong, 2000). Given the results, there is a tendency for the mayonnaise reused bottles still to abhor microorganisms despite the pre-steam step before hot filling.

### Metagenomic analysis of pepper pastes

A full-length 16s metagenomic analysis was carried out for bacterial DNA sequences (100%) to understand the diversity of bacteria in stored pepper samples. Both pastes were colonised mainly by the phyla Proteobacteria and Firmicutes (Figure 1 (a and b). The sample processed with onions had the highest Proteobacteria occurrence (51%), followed by Firmicutes (49%), whereas the sample with no onions had the highest Firmicutes (85%), followed by Proteobacteria (12%). Sequences of organisms that could not be assigned to any known phylum accounted for 0.3% in sample A and 0.2% in ‘B’. Specifically, the two samples yielded bacterial sequences from the Firmicutes, Proteobacteria, and Bacteroidota taxa. In addition to these, the bacterial phyla Actinobacteriota, Acidobacteria, Myxococcota, and Bdellovibrionota were also present in sample B. Mamphogoro et al. (2020) noted that the most abundant sequences from the surfaces of pepper fruits were affiliated with the phylum Proteobacteria, followed by Firmicutes. It could be inferred that the colonies of the raw materials were not wiped out during processing and were not made dormant with the storage method used. Regardless of processing, the core microbiome of the Chinese traditional red pepper paste was also identified and mainly assigned to Proteobacteria and Firmicutes phyla (Li et al., 2016).

**Figure 1:**
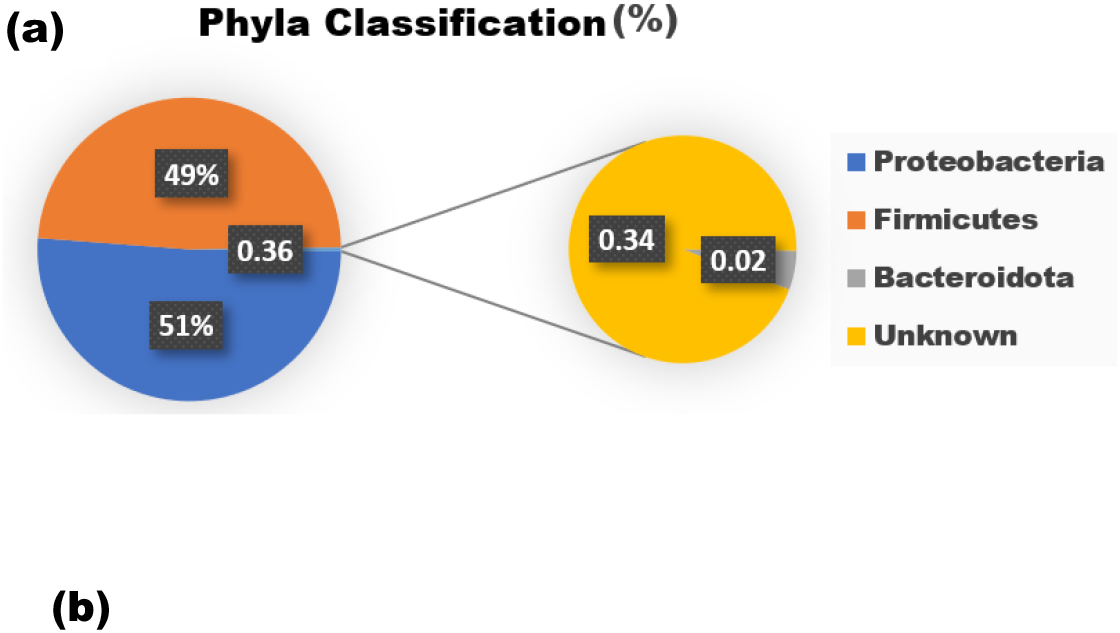
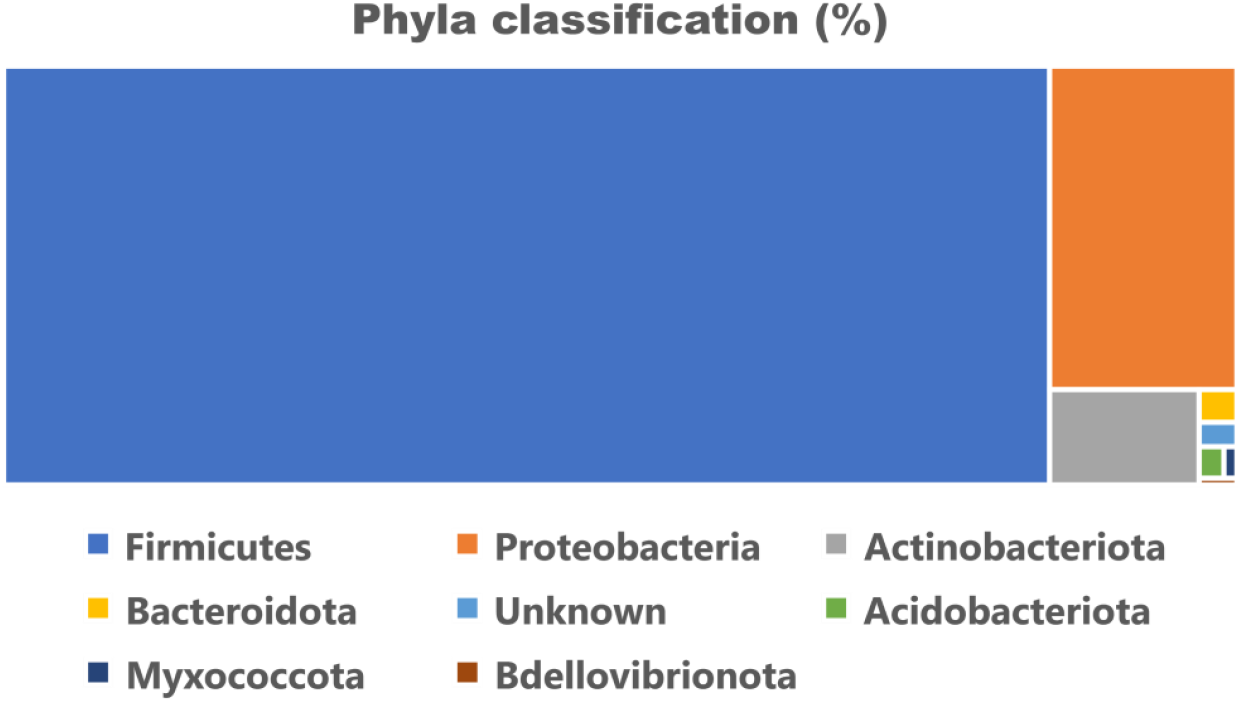
(a.) Cumulative proportion of full-length 16S Ribosomal RNA gene sequences obtained as different bacterial phyla linked with stored pepper paste with onions (Sample A) (total number of bacterial sequences read count = 42,642) (b.) Cumulative proportion of full-length 16S Ribosomal RNA gene sequences obtained as different bacterial phyla linked with stored pepper paste without onions (Sample B) (total number of bacterial sequences read count = 15,313)

The Proteobacteria represent the most legitimately published prokaryotic genera and comprise a significant portion of traditional Gram-negative bacteria. They exhibit remarkable metabolic diversity and are highly significant to biology since they contain the most known Gram-negatives with applications in agriculture, veterinary medicine, industry, and medicine (Kersters et al., 2006). Proteobacteria can survive in various oxic environments since they are often facultatively or obligately anaerobic. Therefore, it is assumed that the Proteobacteria support the stability of the strictly anaerobic microbiota by helping to maintain the anaerobic environment of the GI tract in a state of homeostasis. Also, they are essential in priming the gut for subsequent colonisation by the stringent anaerobes necessary for gut health by absorbing oxygen and reducing redox potential (Moon et al., 2018). However, Proteobacteria can grow in the mucous layer of the stomach that secretes acid, where multiple species of *Helicobacter* have been found raising the pH of their immediate surroundings (Sidebotham et al., 2003). According to a recent investigation, Proteobacteria, despite their relatively lower abundance in the human gut microbiome, account for a large portion of the microbiome functional variance. The implication is that the primary taxonomic sources of microbiota variation (Firmicutes and Bacteroidetes) may not always account for the most significant variation in function (Bradley and Pollard, 2017).

One of the most important external elements influencing the gut microbial architecture is thought to be diet (Leeming et al., 2019). Thus, consuming pepper paste containing a substantial amount of the species could lead to a bloom of Proteobacteria in the gut, resulting in imbalanced gut microbiota, a vital factor determining host health when considering inflammation and metabolism. Numerous studies to date endorse that an abnormal expansion of Proteobacteria would compromise the ability to maintain a balanced gut microbial community, a potential diagnostic marker of dysbiosis and illness risk (Mazmanian et al., 2008). The healthy mammalian gut contains several members of commensal bacterial species belonging to this phylum as its natural gut flora. These commensals seem benign in minor proportion, but they become colitogenic microbes under specific gut environments that can trigger inflammatory responses. Members of the phylum Firmicutes are widely distributed in aquatic and soil settings, where they participate in the recycling and breakdown of organic waste. Several genera are harmful to people, animals, or plants. A few species of the phylum are also helpful to industry in manufacturing enzymes and antibiotics (Seong et al., 2018). Firmicutes have many conserved genes that contribute to the microbiome’s functional redundancy and rank among the most prevalent types of bacteria that make up the human microbiome (Flint et al., 2007).

The findings of the present work demonstrated that the inclusion or non-inclusion of onions in the paste determined the bacteria’s taxonomic structure and diversity (Table 2). Pepper paste with onions (‘A’) had a Gini-Simpson index (or Simpson’s index of diversity) of 0.51, while pepper paste without onions (‘B’) had a 0.18 index. Thus, the higher score in ‘A’ indicates more bacteria diversity than in ‘B’ with respect to even the population proportion of species despite there being more species in ‘B’. The findings showed that a limited group of bacteria thrived in ‘A’ at a reasonably good distribution, but ‘B’ permitted the growth of more species with very poor distribution. Comparing diversity metrics that indicate the number of groups (often species) in an assemblage (richness) or the distribution of those groups within the assemblage (evenness) can be enlightening from a classical ecological perspective. High diversity is correlated with high richness, and an assemblage that is highly dominated (i.e., has poor evenness) is thought to be less diversified than one that is more even. Since the two components of diversity—richness and evenness—calculated from the same samples may exhibit divergent patterns, Simpson’s diversity index evaluates them both (Somerfield et al., 2008).

**Table 2.**
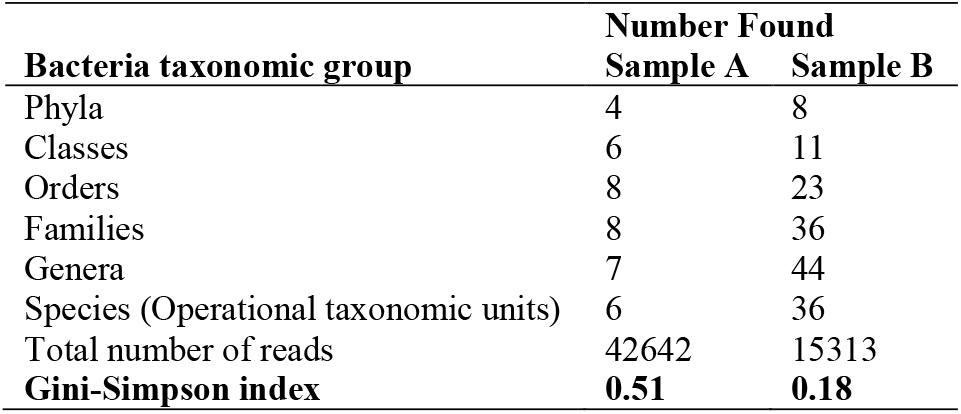
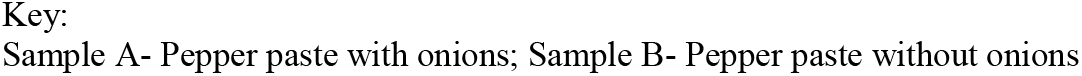
Effect of pepper constituents on the bacterial ecology of stored pepper pastes.

| **Bacteria taxonomic group** | **Number Found** |  |
| --- | --- | --- |
|  | **Sample A** | **Sample B** |
| Phyla | 4 | 8 |
| Classes | 6 | 11 |
| Orders | 8 | 23 |
| Families | 8 | 36 |
| Genera | 7 | 44 |
| Species (Operational taxonomic units) | 6 | 36 |
| Total number of reads | 42642 | 15313 |
| **Gini-Simpson index** | **0.51** | **0.18** |

Sample A- Pepper paste with onions; Sample B- Pepper paste without onions

Sample ‘A’ had 42,642 bacterial sequences belonging to 6 OTUs (Operational taxonomic units) representing bacterial species in 7 Genera, 8 Families, 8 Orders, 6 Classes and 4 phyla. Meanwhile, paste without onions had 15,313 bacterial sequences belonging to 36 OTUs in 44 genera of 36 Families and 23 Orders belonging to 11 Classes and 8 phyla. The study also revealed that sample A is colonised majorly by members of the Classes Gammaproteobacteria (Genus *Proteus;* 50.96% translating to 21,745 read counts) and Bacilli (Genus *Pediococcus*; 48.32% translating to 20,713 read counts) (Table 3). Taxonomic and systematic groupings of the 16S rRNA gene sequences belonging to the phylum Firmicutes were majorly represented by members of the Class Bacilli and genus *Bacillus* (83.98% translating to 12,856 read counts) in sample B. The least occurring Class in ‘B’ is Acidobacteria, represented by the genus *Acidibacter* at 0.16% with 25 read counts. Alphaproteobacteria and Actinobacteria are major classes in ‘B’ that are absent in ‘A’. The Class Gammaproteobacteria occurring in ‘B’ spanned across 13 genera: *Proteus, Providencia, Klebsiella, Escherichia, Enterobacter, Serratia*; *Pseudomonas, Acinetobacter, Psychrobacter, Azotobacter, Oligella, Halomonas, Stenotrophomonas*.

**Table 3.** Relative percentage occurrence of Bacterial genera of classes and orders associated with stored pepper paste.

| Class | Order | Genus | Sample A (%) | Sample B (%) |
| --- | --- | --- | --- | --- |
| Bacilli | Bacillales | *Bacillus* | 0.05 | 83.98 |
|  |  | *Lysinibacillus* | ND | 0.13 |
|  | Lactobacillales | *Pediococcus* | 48.32 | 0.02 |
|  |  | *Lactococcus* | 0.07 | ND |
|  |  | *Enterococcus* | 0.02 | 0.14 |
|  |  | *Vagococcus* | ND | 0.11 |
| Alphaproteobacteria | Sphingomonadales | *Sphingomonas* | ND | 3.57 |
|  | Rhizobiales | *Methylobacterium* | ND | 2.00 |
|  |  | *Bhargavaea* | ND | 0.25 |
|  |  | *Brevundimonas* | ND | 0.18 |
|  | Rhodospirillales | *Rhodoplanes* | ND | 0.10 |
|  |  | *Azospira* | ND | 0.02 |
| Gammaproteobacteria | Enterobacterales | *Proteus* | 50.96 | 1.70 |
|  |  | *Providencia* | ND | 0.35 |
|  |  | *Klebsiella* | ND | 0.29 |
|  |  | *Escherichia* | ND | 0.24 |
|  |  | *Enterobacter* | ND | 0.16 |
|  |  | *Serratia* | ND | 0.11 |
|  | Pseudomonadales | *Pseudomonas* | ND | 0.85 |
|  |  | *Acinetobacter* | ND | 0.33 |
|  |  | *Psychrobacter* | ND | 0.24 |
|  |  | *Azotobacter* | ND | 0.03 |
|  | Oceanospirillales | *Oligella* | ND | 0.12 |
|  |  | *Halomonas* | ND | 0.08 |
|  | Xanthomonadales | *Stenotrophomonas* | ND | 0.07 |
| Actinobacteria | Propionibacterales | *Cutibacterium* | ND | 1.69 |
|  |  | *Nocardioides* | ND | 0.02 |
|  | Micrococcales | *Arthrobacter* | ND | 0.57 |
|  |  | *Leucobacter* | ND | 0.10 |
|  |  | *Kocuria* | ND | 0.08 |
|  |  | *Cellulomonas* | ND | 0.03 |
|  |  | *Leifsonia* | ND | 0.03 |
|  | Corynebacterales | *Corynebacterium* | ND | 0.06 |
| Bacteroidetes | Flavobacteriales | *Myroides* | ND | 0.18 |
| Acidobacteria | Acidobacteriales | *Acidibacter* | ND | 0.16 |
| Betaproteobacteria | Burkholderiales | *Alcaligenes* | ND | 0.16 |
|  |  | *Enhydrobacter* | ND | 0.03 |
| Bacteroidia | Bacteroidales | *Sediminibacterium* | ND | 0.06 |
| Negativicutes | Acidaminococcales | *Phascolarctobacterium* | ND | 0.05 |
| Proteobacteria | Myxococcales | *Archangium* | ND | 0.05 |
| Clostridia | Clostridiales | *Faecalibacterium* | 0.01 | ND |
| Unknown | Unknown | Unknown | 0.56 | 1.18 |
|  |  | uncultured | ND | 0.37 |
|  |  | RB41 | ND | 0.05 |
|  |  | 0319 | ND | 0.05 |
|  |  | *Amphiplicatus* | ND | 0.01 |

Sample A- Pepper paste with onions; Sample B- Pepper paste without onions; ND- Not detected

Gammaproteobacteria is the largest group in Proteobacteria and encompasses several well-known chemoorganotrophic bacterial orders, including Enterobacterales and Pseudomonadales. Several significant human, animal and plant pathogens are present in the Class. The core of the γ-class is made up of well-known enterics, among others (Kersters et al., 2006). Several members of the digestive tract of homoiothermic animals, including the extensively researched *Escherichia coli*, are members of the Enterobacterales Order. *E. coli*, an excellent indicator organism, effectively detects faecal pollution in water. Fortunately, *E. coli* was not detected in the pastes; uncultured *Escherichia* was found in the sample with no onions at 0.24%, amounting to 37 bacterial sequences in the genus. Plant pathogenic bacteria belonging to the *Escherichia* genus could have been part of the raw materials’ microbiota that were not eliminated by the production method. *Proteus*, the highest occurring genus in the sample with onions (‘A’) is a member of the Enterobacteriaceae family. They are found in multiple environmental habitats, including higher organisms, soil, and water. It is hypothesised that intestines are a reservoir of these organisms (Zilberstein et al., 2007) where they act as opportunistic human pathogens, commensals, symbionts or their presence handled as a carrier status (Drzewiecka, 2016). *Proteus* is an undesired element of intestinal microflora, as the bacteria may become a causative agent of diarrhoea (Müller, 1989). Cross-contamination from production handlers may have caused the presence of Proteus in both samples, possibly due to poor hygiene practices (Shojaei et al., 2006). It can also be deduced that this group of bacteria is not affected by the antimicrobial effect of onions and thus proliferates a lot in sample A. Their presence in the pepper pastes also indicates deterioration, as *Proteus* can tolerate or utilise polluting compounds (Drzewiecka, 2016).

The Bacilli Class is widely present in various natural environments and is extensively studied in soil, saline water, plants, animals, and air. The *Bacillus* genus showcases a wide array of phenotypic diversity, including the use of uncommon terminal electron acceptors such as arsenic or selenium (Blum’ et al., 1998). Other members of the genus are tolerant of high temperatures, extreme salinity, acidic conditions, and the immune systems of many animals. They are distinguished by cells that develop aerobically and produce dormant endospores. (Maughan and Van der Auwera, 2011). The raw material microbiota is likely to have a significant quantity of *Bacillus*, specifically their spores, which is why the sample lacking onions (designated as “B”) has the highest concentration of *Bacillus*.

The majority (90.35%, equating to 13,829 read counts) of the bacteria in the sample with no onions (‘B’) could not be assigned to any taxonomic group at the species level (Table 4). *Sphingomonas echinoides* (3.42%) is the most occurring species, and *Cutibacterium sp*. (0.01) is the least in ‘B’. If the former is the highest occurring identified species, it could be inferred that most of the unknown species in ‘B’ would belong to the genus *Bacillus. Sphingomonas* is found in a wide range of habitats and has been isolated from soils, even those contaminated with pollutants, plant roots, and water distribution systems. Some species cause disease, some are antagonistic to other microbes, others aid the extraction of rare earth elements, some serves as biocatalyst for bioremediation, and some produce highly beneficial phytohormones (Sorouri et al., 2023). *Sphingomonas echinoides* was first isolated as a plate contaminant (Shin et al., 2012). *Cutibacterium sp*., formerly called *Propionibacterium sp*., is considered commensal skin bacteria and contaminants that are typically non-pathogenic (Corvec, 2018). To prevent these organisms from pepper paste, a series of stages of bacterial control must be incorporated into the manufacturing procedure. *Proteus mirabilis* (42.92%) dominated Sample A, followed by *Pediococcus pentosaceus* (40.73%), and the least occurring species is *Lactococcus garvieae* (0.04%). *P. mirabilis* causes the majority of *Proteus* community-acquired infections. Wang et al. (2010) documented a case of *P. mirabilis*-related food poisoning that occurred in a Beijing, China restaurant. Additional research supports that unclean hands could significantly transmit *Proteus species* from faeces to hands and mouths.

**Table 4.**
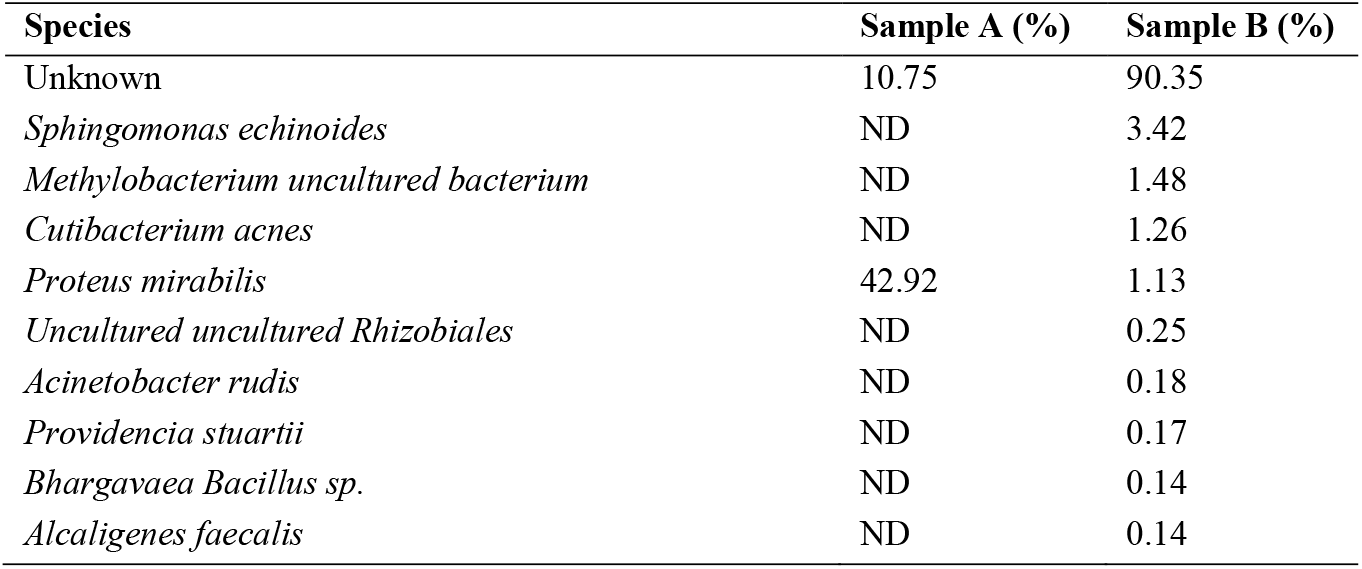
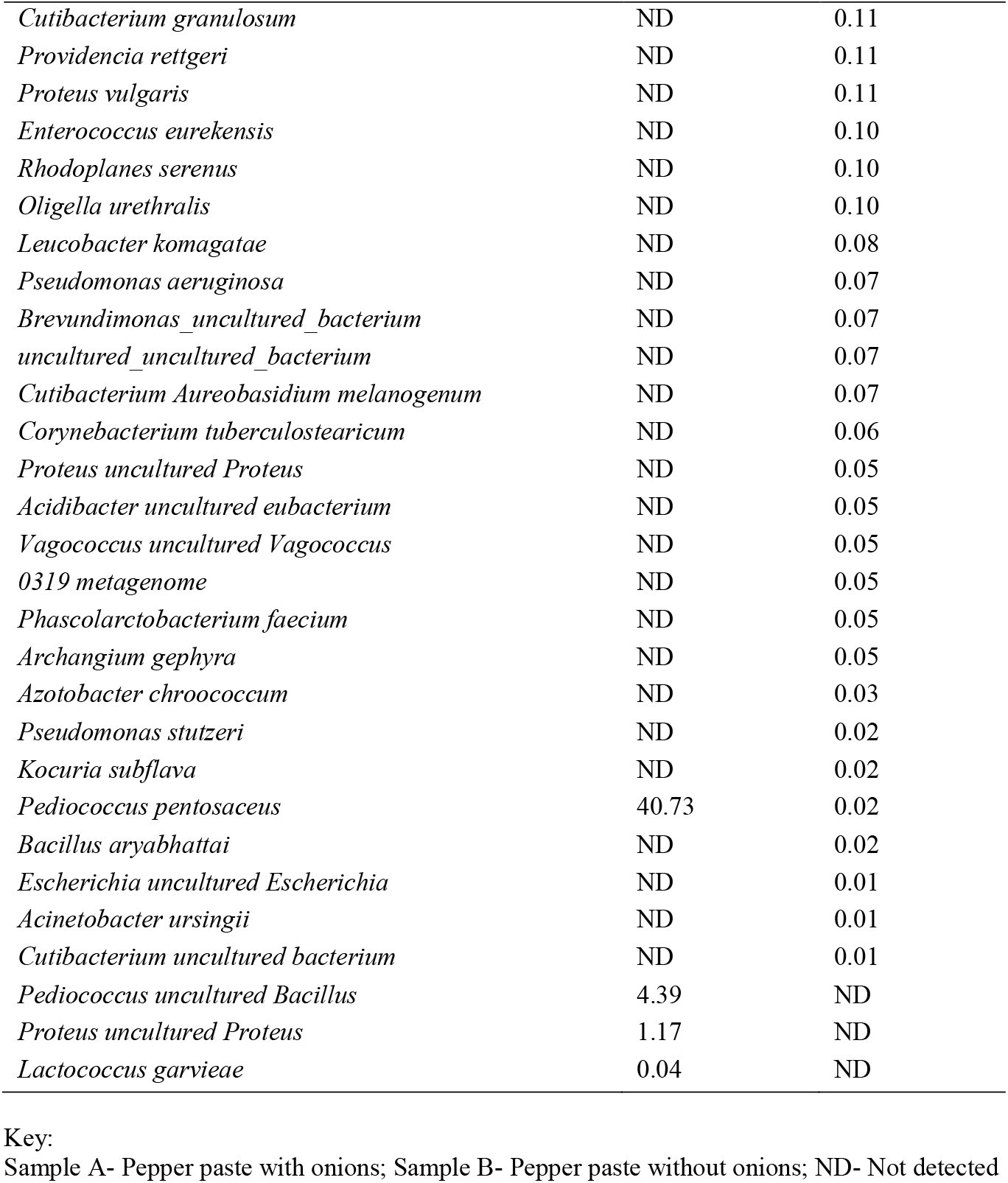
Relative percentage occurrence of bacterial species associated with stored pepper paste.

*P. mirabilis* was discovered to colonise hand skin between the nail plate and nail fold in motor mechanics by Qadripur (2001).

The presence of *Pediococcus pentosaceus* in sample A suggests its antagonistic properties against numerous species found in ‘B’. *P. pentosaceus* occurs in food as native microflora and in fermentations as starters. It has an antimicrobial effect, providing a natural means of preservation. Hence, the food industry uses the bacteria’s cultures or products as bio- preservatives (Papagianni and Anastasiadou, 2009). European Food Safety Authority (EFSA) also found a variant of *P. pentosaceus* that is appropriate for the qualified presumption of safety (QPS) approach (Bampidis et al., 2022). Since the first human case was reported in 1991, *L. garvieae*, the least occurring species in ‘A’, has become known as a newly discovered opportunistic human disease. Among other foods, the pathogen has also been isolated from vegetables and cereals (Masudi et al., 2023).

### Chemical composition of stored pepper pastes

There is no direct correlation between the two samples’ titratable acidity (TA) and pH (Table 5). In the sample with onions (‘A’), the TA decreased significantly (p < 0.05) within 30 days of storage (0.43 to 0.41%) and then increased consistently. Meanwhile, the pH decreased steadily from the day of production (5.11), came to 4.20 on the 60^th^ day of storage, and then increased (4.68). Organic acids formed due to metabolic activities of the microorganisms in the pepper paste would have reduced the pH initially. Bozkurt and Erkmen (2004) observed the same trend and opined that organic acids would break down as the storage period increases and specific compounds might form. As a result, the pastes’ pH levels rose. In ‘B’, the TA values swing (p < 0.05) by first rising significantly (0.47 to 0.50%) and then falling (0.45%). The pH values were reduced by day 30 (4.90 to 4.13) and rose by the 60^th^ day (4.36). Generally, ‘B’ maintained acid pH. Throughout the storage period, the pepper paste had high dry matter (DM) losses (p < 0.05). By the 30^th^ day of storage, the amount of DM in both samples had dropped by an average of 52% (‘A’: 24.43 to11.72%; ‘B’: 28.53 to 13.42%). It stayed at this level in sample A until the 90^th^ day, when it dropped again (10.17%). However, in ‘B,’ it steadily decreased and held its value from the 60^th^ (11.31%) to the 90^th^ (11.26%) day. Dry matter loss from pastes also translates to a loss in the ash content of the samples and a loss in the pastes’ quality (Akakpo et al., 2020). During the storage periods, there were significant losses in the contents of ash and total carotenoids in both samples, with a consistent decline in value.

**Table 5.** Percentage Titratable acidity, Dry matter, Ash content, Carotenoids and pH of pepper pastes over days of storage.

| Sample | Parameter | 0 | 30 | 60 | 90 |
| --- | --- | --- | --- | --- | --- |
| **A** | Titratable Acidity (%) | 0.43±0.03<sup>ab</sup> | 0.41±0.01<sup>a</sup> | 0.42±0.03<sup>ab</sup> | 0.45±0.01<sup>b</sup> |
|  | Dry matter (%) | 24.43±1.21<sup>b</sup> | 11.72±0.11<sup>ab</sup> | 11.70±0.55<sup>ab</sup> | 10.17±0.54<sup>a</sup> |
|  | Ash content (%) | 2.11±0.03<sup>c</sup> | 0.81±0.01<sup>b</sup> | 0.77±0.00<sup>a</sup> | 0.77±0.01<sup>a</sup> |
|  | Total carotenoids (mg g<sup>-1</sup>) | 2.11±0.03<sup>d</sup> | 0.43±0.01<sup>c</sup> | 0.38±0.01<sup>b</sup> | 0.23±0.00<sup>a</sup> |
|  | pH | 5.11±0.03<sup>d</sup> | 4.34±0.01<sup>b</sup> | 4.20±0.00<sup>a</sup> | 4.68±0.02<sup>c</sup> |
| **B** | Titratable Acidity (%) | 0.47±0.02<sup>ab</sup> | 0.50±0.01<sup>b</sup> | 0.45±0.00<sup>a</sup> | 0.45±0.03<sup>a</sup> |
|  | Dry matter (%) | 28.53±1.21<sup>c</sup> | 13.42±0.11<sup>b</sup> | 11.31±0.53<sup>a</sup> | 11.26±0.13<sup>a</sup> |
|  | Ash content (%) | 1.88±0.03<sup>c</sup> | 1.21±0.01<sup>b</sup> | 1.19±0.02<sup>b</sup> | 0.19±0.00<sup>a</sup> |
|  | Total carotenoids (mg g<sup>-1</sup>) | 1.65±0.02<sup>c</sup> | 0.44±0.02<sup>b</sup> | 0.25±0.00<sup>a</sup> | 0.23±0.03<sup>a</sup> |
|  | pH | 4.90±0.03<sup>c</sup> | 4.13±0.01<sup>a</sup> | 4.36±0.02<sup>b</sup> | 4.37±0.00<sup>b</sup> |

Sample A- Pepper paste with onions; Sample B- Pepper paste without onions

### Physical assessment of stored pepper pastes

Three onion-free samples showed signs of mould growth by 30 days. By day 60, mould had developed in five of the onion-containing samples (“A”) and nine of the onion-free samples (“B”). Some of the latter also gave off a strong, yeasty, and mouldy odour. By day 90, three of the “A” samples and eight of the “B” samples showed visible mould growth. Sample A appeared to be holding up well at day 120, while no “B” samples remained free from spoilage. Some of the spoiled pepper pastes had white, fuzzy patches and black spots, while others were still slimy to the touch or had a pungent odour. Despite some cottage producers claiming that their paste could last 9 to 12 months without physical symptoms of spoilage, a few individuals who make pepper paste for personal consumption too reported experiencing spoilage within 30 days.

## Conclusion

Findings from this work have provided insights into the factors that affect the quality and safety of stored pepper pastes using microbiological and chemical indicators. Metagenomic analysis revealed the microbial communities in the products as related to the inclusion or non-inclusion of onions and their implications for quality and safety. The microbiota of stored pepper paste basically reflects the microbial communities in the soil, human hands, skin, and air. Identifying the microorganisms gave an insight into those responsible for spoilage, food poisoning or gut microbiota imbalance. This study found that pepper paste with onions has a shelf-life of 60 days and the one without onions 30 days, regardless of whether there are visible symptoms of deterioration. It would be interesting to have standard organisations issue guidelines for producers that make food for domestic consumption regarding minimum reasonable hygienic practice requirements. Part of this should include controls to be used during processing and packing cottage or kitchen food products to ensure that they meet quality.

## REFERENCES

Akakpo, D. B., de Boer, I. J. M., Adjei-Nsiah, S., Duncan, A. J., Giller, K. E., & Oosting, S. J. (2020). Evaluating the effects of storage conditions on dry matter loss and nutritional quality of grain legume fodders in West Africa. Animal Feed Science and Technology, 262. 10.1016/j.anifeedsci.2020.114419

AOAC. (2000). Official methods of analysis of AOAC. International 17th edition; Gaithersburg, MD, USA Association of Analytical Communities.

Aşkin, U. R. (2018). Preservation of sweet red pepper paste quality: Effect of packing material, ozone gas and protective agent use. Food Science and Technology (Brazil), 38(4), 698–703. 10.1590/1678-457x.13917

Bampidis, V., Azimonti, G., Bastos, M. de L., Christensen, H., Dusemund, B., Fašmon Durjava, M., Kouba, M., López-Alonso, M., López Puente, S., Marcon, F., Mayo, B., Pechová, A., Petkova, M., Ramos, F., Sanz, Y., Villa, R. E., Woutersen, R., Saarela, M., Anguita, M., … Revez, J. (2022). Safety and efficacy of a feed additive consisting of Pediococcus pentosaceus DSM 32292 for all animal species (Marigot Ltd t/a Celtic Sea Minerals). EFSA Journal, 20(7). 10.2903/j.efsa.2022.7426

Banwart, G. J. (1989). Sources of microorganisms. In: Basic Food Microbiology. Springer, Boston, MA. 10.1007/978-1-4684-6453-5_5

Blum’, J. S., Burns, A., Buzzelli, B. J., Stolz’, J. F., & Oremland’, R. S. (1998). Bacillus arsenicoselenatis, sp. noy., and Bacillus selenitireducens, sp. noy.: two haloalkaliphiles from Mono Lake, California that respire oxyanions of selenium and arsenic. In Arch Microbiol (Vol. 171).

Bozkurt, H., & Erkmen, O. (2004). Effects of production techniques on the quality of hot pepper paste. Journal of Food Engineering, 64(2), 173–178. 10.1016/j.jfoodeng.2003.09.028

Bradley, P. H., & Pollard, K. S. (2017). Proteobacteria explain significant functional variability in the human gut microbiome. Microbiome, 5(1). 10.1186/s40168-017-0244-z

Chakraborty, A. J., Uddin, T. M., Matin Zidan, B. M. R., Mitra, S., Das, R., Nainu, F., Dhama, K., Roy, A., Hossain, M. J., Khusro, A., & Emran, T. Bin. (2022). Allium cepa: A Treasure of Bioactive Phytochemicals with Prospective Health Benefits. In Evidence-based Complementary and Alternative Medicine (Vol. 2022). Hindawi Limited. 10.1155/2022/4586318

Chauhan, A., & Jindal, T. (2020). Microbiological Methods for Environment, Food and Pharmaceutical Analysis. In Microbiological Methods for Environment, Food and Pharmaceutical Analysis. Springer International Publishing. 10.1007/978-3-030-52024-3

Chitravathi, K., Chauhan, O. P., & Raju, P. S. (2015). Influence of modified atmosphere packaging on shelf-life of green chillies (Capsicum annuum L.). Food Packaging and Shelf Life, 4, 1–9. 10.1016/J.FPSL.2015.02.001

Corvec, S. (2018). Clinical and Biological Features of Cutibacterium (Formerly Propionibacterium) avidum, an Underrecognized Microorganism. https://journals.asm.org/journal/cmr

Drzewiecka, D. (2016). Significance and Roles of Proteus spp. Bacteria in Natural Environments. Microbial Ecology, 72(4), 741–758. 10.1007/s00248-015-0720-6

Etebu, E., Torunana, J. M. A., & Parker, M. (2018). Metagenomic analysis of bacterial community associated with postharvest Irvingia species fruit wastes. Microbiology Research International, 6(2), 7–15. 10.30918/MRI.62.18.014

Falola, A., Mukaila, R., Uddin II, R. O., Ajewole, C. O., & Gbadebo, W. (2023). Postharvest losses in onion: causes and determinants. Kahramanmaraş Sütçü Imam Üniversitesi Tarim ve Doğa Dergisi, 26(2), 346–354. 10.18016/ksutarimdoga.vi.1091225

Fawole, A. O. (2019). Selection of lactic acid bacteria for use as starter cultures in lafun production and their impact on product quality and safety [Doctoral thesis, University of Reading]. British Library, the National Library of the United Kingdom database (ETOS)

Flint, H. J., Duncan, S. H., Scott, K. P., & Louis, P. (2007). Interactions and competition within the microbial community of the human colon: Links between diet and health: Minireview. In Environmental Microbiology (Vol. 9, Issue 5, pp. 1101–1111). 10.1111/j.1462-2920.2007.01281.x

Kabrah, A., Faidah, H., & Ashshi, A. (2016). Antibacterial Effect of Onion Developing a 3D human Bioreactor for stem cell studies View project Affect of Onion on MSSA and MRSA isolate from healthy people View project. 10.21276/sjams.2016.4.11.53

Kersters, K., De Vos, P., Gillis, M., Swings, J., Vandamme, P., & Stackebrandt, E. (2006). Introduction to the Proteobacteria. In The Prokaryotes (pp. 3–37). Springer New York. 10.1007/0-387-30745-1_1

Lau, O.-W., & Wong, S.-K. (2000). Contamination in food from packaging material. In Journal of Chromatography A (Vol. 882). https://www.elsevier.com/locate/chroma

Leeming, E. R., Johnson, A. J., Spector, T. D., & Le Roy, C. I. (2019). Effect of Diet on the Gut Microbiota: Rethinking Intervention Duration. Nutrients, 11(12), 2862. 10.3390/nu11122862

Li, Z., Dong, L., Huang, Q., & Wang, X. (2016). Bacterial communities and volatile compounds in Doubanjiang, a Chinese traditional red pepper paste. Journal of Applied Microbiology, 120(6), 1585–1594. 10.1111/jam.13130

Mamphogoro, T. P., Maboko, M. M., Babalola, O. O., & Aiyegoro, O. A. (2020). Bacterial communities associated with the surface of fresh sweet pepper (Capsicum annuum) and their potential as biocontrol. Scientific Reports, 10(1). 10.1038/s41598-020-65587-9

Masudi, B., Litvinchuk, T., & Byrd, J. (2023). Lactococcus garvieae in Rural Alabama: A Case Report. In Cureus (Vol. 15, Issue 5). 10.7759/cureus.39560

Maughan, H., & Van der Auwera, G. (2011). Bacillus taxonomy in the genomic era finds phenotypes to be essential though often misleading. In Infection, Genetics and Evolution (Vol. 11, Issue 5, pp. 789–797). 10.1016/j.meegid.2011.02.001

Mazmanian, S. K., Round, J. L., & Kasper, D. L. (2008). A microbial symbiosis factor prevents intestinal inflammatory disease. Nature, 453(7195), 620–625. 10.1038/nature07008

Moon, C. D., Young, W., Maclean, P. H., Cookson, A. L., & Bermingham, E. N. (2018). Metagenomic insights into the roles of Proteobacteria in the gastrointestinal microbiomes of healthy dogs and cats. In MicrobiologyOpen (Vol. 7, Issue 5). Blackwell Publishing Ltd. 10.1002/mbo3.677

Müller, H. E. (1989). The Role of Proteae in Diarrhea. Zentralblatt Fur Bakteriologie, 272(1), 30–35. 10.1016/S0934-8840(89)80089-1

Oyawoye, O. M., Olotu, T. M., Nzekwe, S. C., Idowu, J. A., Abdullahi, T. A., Babatunde, S. O., Ridwan, I. A., Batiha, G. E., Idowu, N., Alorabi, M., & Faidah, H. (2022). Antioxidant potential and antibacterial activities of Allium cepa (onion) and Allium sativum (garlic) against the multidrug resistance bacteria. Bulletin of the National Research Centre, 46(1). 10.1186/s42269-022-00908-8

Papagianni, M., & Anastasiadou, S. (2009). Pediocins: The bacteriocins of Pediococci. Sources, production, properties and applications. In Microbial Cell Factories (Vol. 8, Issue 1). BioMed Central Ltd. 10.1186/1475-2859-8-3

Qadripur, S. A., Schauder, S., & Schwartz, P. (2001). Black nails from Proteus mirabilis colonisation. In Der Hautarzt (Vol. 52, pp. 658–661).

Quigley, L., O’Sullivan, O., Beresford, T. P., Ross, R. P., Fitzgerald, G. F., & Cotter, P. D. (2012). High-throughput sequencing for detection of subpopulations of bacteria not previously associated with artisanal cheeses. Applied and Environmental Microbiology, 78(16), 5717–5723. 10.1128/AEM.00918-12

Russell, A. D. (2003). Lethal effects of heat on bacterial physiology and structure. Science Progress, 86(2), 115–137. https://www.scilet.com

Santas, J., Almajano, M. P., & Carbó, R. (2010). Antimicrobial and antioxidant activity of crude onion (Allium cepa, L.) extracts. International Journal of Food Science and Technology, 45(2), 403–409. 10.1111/j.1365-2621.2009.02169.x

Seong, C. N., Kang, J. W., Lee, J. H., Seo, S. Y., Woo, J. J., Park, C., Bae, K. S., & Kim, M. S. (2018). Taxonomic hierarchy of the phylum Firmicutes and novel Firmicutes species originated from various environments in Korea. In Journal of Microbiology (Vol. 56, Issue 1). Microbiological Society of Korea. 10.1007/s12275-018-7318-x

Shin, S. C., Kim, S. J., Ahn, D. H., Lee, J. K., & Park, H. (2012). Draft genome sequence of Sphingomonas echinoides ATCC 14820. Journal of Bacteriology, 194(7), 1843. 10.1128/JB.00046-12

Shojaei, H., Shooshtaripoor, J., & Amiri, M. (2006). Efficacy of simple hand-washing in reduction of microbial hand contamination of Iranian food handlers. Food Research International, 39(5), 525–529. 10.1016/j.foodres.2005.10.007

Sidebotham, R. L., Worku, M. L., Karim, Q. N., Dhir, N. K., & Hugh, B. J. (2003). How Helicobacter pylori urease may affect external pH and influence growth and motility in the mucus environment: evidence from in-vitro studies. European Journal of Gastroenterology & Hepatology, 15, 395–401. 10.1097/01.meg.0000050029.34359.98

Somerfield, P. J., Clarke, K. R., & Warwick, R. M. (2008). Simpson Index. In Encyclopedia of Ecology, Five-Volume Set (pp. 3252–3255). Elsevier Inc. 10.1016/B978-008045405-4.00133-6

Sorouri, B., Rodriguez, C. I., Gaut, B. S., & Allison, S. D. (2023). Variation in Sphingomonas traits across habitats and phylogenetic clades. Frontiers in Microbiology, 14. 10.3389/fmicb.2023.1146165

Wang, Y., Zhang, S., Yu, J., Zhang, H., Yuan, Z., Sun, Y., Zhang, L., Zhu, Y., & Song, H. (2010). An outbreak of Proteus mirabilis food poisoning associated with eating stewed pork balls in brown sauce, Beijing. Food Control, 21(3), 302–305. 10.1016/j.foodcont.2009.06.009

Zilberstein, B., Quintanilha, A. G., Santos, M. A. A., Pajecki, D., Moura, E. G., Alves, P. R. A., Maluf Filho, F., De Souza, J. A. U., & Gama-Rodrigues, J. (2007). Digestive tract microbiota in healthy volunteers. Clinics, 62(1), 47–54. 10.1590/S1807-59322007000100008

